# Role of Thioredoxin Reductase (TrxB) in Oxidative Stress Response of *Francisella tularensis* Live Vaccine Strain

**DOI:** 10.1101/2025.04.27.650861

**Authors:** Matthew Higgs, Zhuo Ma, Anthony Centone, Chandra Shekhar Bakshi, Meenakshi Malik

## Abstract

*Francisella tularensis* is an important human pathogen responsible for causing tularemia in the Northern Hemisphere. *Francisella* has been developed as a biological weapon in the past due to its extremely high virulence. *F. tularensis* is a Gram-negative, intracellular pathogen that primarily infects macrophages. To counteract the reactive oxygen and nitrogen species (ROS/RNS) produced by macrophages in response to infection, *F. tularensis* encodes a repertoire of antioxidant enzymes. Among these, the thioredoxin system is critical for maintaining cellular redox homeostasis by regulating the balance between oxidation and reduction within bacterial cells. This system includes thioredoxins, thioredoxin reductase, and NADPH. Despite its potential importance, the thioredoxin system of *F. tularensis* remains understudied. *F. tularensis* possesses two thioredoxin genes, *trxA1* (*FTL_0611*) and *trxA2* (*FTL_1224*), and a single thioredoxin reductase gene, *trxB* (*FTL_1571*). In this study, we characterized the role of *trxB* in oxidative stress resistance. Our findings demonstrate that *trxB* is essential for oxidative stress resistance in *F. tularensis* and that its loss increases susceptibility to several antibiotics. However, unlike in other bacterial species, TrxB in *F. tularensis* is not a functional target of the gold-containing antimicrobial agent auranofin. We also show that OxyR, the master regulator of oxidative stress responses, directly controls *trxB* expression under oxidative stress conditions. Furthermore, TrxB contributes to intramacrophage survival by enabling the bacterium to withstand ROS-induced oxidative stress. Collectively, this study highlights a critical, previously uncharacterized antioxidant defense mechanism in *F. tularensis*, highlighting its importance in oxidative stress resistance and intramacrophage survival.

## Introduction

*Francisella tularensis* is an important human pathogen responsible for causing a fatal disease known as tularemia in the northern hemisphere. *F. tularensis* has been developed as a biological weapon in the past because of its extremely high virulence. Based on its potential use as a bioterror agent, it is now classified as a Tier 1 category A select agent by the CDC (1, 2). The human virulent *F. tularensis* strains are classified under *F. tularensis* subsp. *tularensis* (type A) and *F. tularensis* subsp. *holarctica* (type B) (3). The highly virulent and category A select agent, *F. tularensis* SchuS4 strain, belongs to *F. tularensis* subsp. *tularensis,* while the live vaccine strain (LVS) is a derivative of *F. tularensis* subsp. *holarctica* developed in the USA from the Russian strain S15 (4). The clinical presentation of tularemia depends on the route, dose, and infecting strain of *F. tularensis*. The ulceroglandular, oculoglandular, or typhoidal forms of tularemia are not fatal. Pneumonic tularemia is a highly acute and fatal form of the disease (5–7). Tularemia is a notifiable disease in the USA. Naturally occurring tularemia is reported from all the states of the USA, except Hawaii (8, 9). The incidence of tularemia has been on the rise in the past few years in the South-Central, the Pacific Northwest, and Massachusetts, including Martha’s Vineyard (10, 11). No vaccine is currently available for tularemia prophylaxis.

1. *F. tularensis* is a Gram-negative intracellular pathogen that primarily infects macrophages. *F. tularensis* has a unique intramacrophage lifestyle which includes a transient phagosomal phase, followed by its escape into the cytosol, where bacterial replication occurs. *F. tularensis* is exposed to oxidative stress at both intracellular locations (12). To resist oxidative and nitrosative stresses posed by reactive oxygen and nitrogen species (ROS and RNS) produced by the infected macrophages, *F. tularensis* encodes a repertoire of antioxidant enzymes to neutralize ROS and RNS. Studies using the point mutant of superoxide dismutase B (*sodB*) gene and deletion mutants of *sodC* (Δ*sodC* and *sodB*Δ*sodC*), alkylhydroperoxy reductase (Δ*ahpC*), catalase gene deletion mutant (Δ*katG*), and oxidative stress transcriptional regulator *oxyR* (Δ*oxyR*) genes have shown that all the single-gene deletion mutants in antioxidant enzyme genes of *F. tularensis* LVS are highly sensitive towards ROS/RNS and are attenuated for virulence in mice, indicating that antioxidant enzymes of *F. tularensis* LVS serve as essential virulence factors (13–17). A transmembrane component of the major facilitator superfamily type of efflux pump, EmrA1 confers resistance to oxidants by secreting antioxidant enzymes SodB and KatG (18). Additionally, the outer membrane component SilC of the Emr multidrug efflux pump also confers resistance against oxidative stress (19). The stringent response in *F. tularensis* LVS governs the oxidative stress response by regulating the expression of genes encoded on the Francisella pathogenicity island, transcriptional regulators, and antioxidant enzyme genes (20). A transcriptional regulator belonging to the AraC/XylS family also modulates the oxidative stress response of *F. tularensis* LVS (21).

The thioredoxin system of *F. tularensis,* required to mitigate oxidative stress encountered during its intracellular lifecycle, remains understudied. The thioredoxin system is essential for cellular redox regulation and maintaining the balance between oxidation and reduction within bacterial cells. It includes thioredoxins, thioredoxin reductase, and NADPH. This system protects bacteria from oxidative stress and regulates DNA synthesis, repair, and apoptosis (22). Thioredoxins are small proteins that act as antioxidants, reducing other proteins through cysteine thiol-disulfide exchange. Thioredoxin reductase regenerates thioredoxins to their reduced form using electrons from NADPH. This system in *Francisella* comprises two thioredoxins, TrxA1 and TrxA2, and a single thioredoxin reductase (TrxR) annotated as TrxB. TrxA1 regulates the oxidative stress response by modulating the expression of the master regulator, OxyR. However, unlike TrxA2, TrxA1 is essential for resisting oxidative stress and survival within macrophages, highlighting its unique mechanism of oxidative stress response regulation, compared to other bacterial thioredoxins (23). In this study, we characterized the role of TxrB of *F. tularensis* in the oxidative stress resistance of *F. tularensis*.

## Results

### Confirmation of *trxB* gene deletion, transcomplementation, and bioinformatic analysis

The thioredoxin reductase (TrxR) is annotated as TrxB in the *Francisella* genome sequence. The *trxB* (*FTL_1571*) gene in *F. tularensis* LVS is 951 base pairs long and encodes a protein of 316 amino acids. The amino acid sequences of TrxB of *F. tularensis* LVS show 98.73% homology to TrxB (*FTT_0489c*) of *F. tularensis* SchuS4 and 99.37% homology to TrxB (*FTN_0580*) of *F. novicida.* Amino acid sequence alignment of TrxB of *Francisella* with *H. pylori*, *M. tuberculosis*, *P. aeruginosa*, *V. cholerae*, *Y. pestis*, *E. coli*, and *S.* Typhi revealed four conserved motifs essential for protein functions (**Fig. S1A**). The GXGXXA motif, which serves as a nucleotide-binding domain that enables the binding of the NADPH cofactor, is critical for the reductase activity of the TrxB (24). The conserved CXXC motif (25), located within the active site, facilitates electron transfer from NADPH to thioredoxin. The presence of cysteine residues at both ends of this motif is particularly important for forming disulfide bonds during redox reactions. The GXGXXA motif is required to maintain the structural integrity of the NADPH-binding pocket, ensuring an efficient cofactor interaction. The HRRXXXR motif is involved in protein-protein interactions and contributes to the specificity and functional activity of the TrxB. The conservation of these functional motifs across various bacterial species signifies their conserved role in TrxRs in maintaining redox homeostasis and resistance against oxidative stress.

The confirmation of deletion of the *trxB* gene in the Δ*trxB* mutant was confirmed by PCR using primers internal to the *trxB* gene. The absence of the *trxB-*specific amplification product, but the presence of a product corresponding to the *sodB* gene of *F. tularensis* that was used as an internal control, confirmed the deletion of the *trxB* gene (**Fig. S1B and C**). Amplification of a *trxB-*specific fragment in the transcomplemented strain confirmed the transcomplementation. Since the *trxB* gene of *F. tularensis* LVS is encoded as an operon, its deletion may impact the expression of genes downstream of it by altering the frame of reading. The clean *in-frame* deletion of the *trxB* gene in the Δ*trxB* mutant was confirmed by DNA sequencing (data not shown).

### TrxB contributes to the oxidative stress resistance of *F. tularensis*

The growth of *F. tularensis* LVS, the Δ*trxB* mutant, and the transcomplemented strain was assessed under untreated and H_2_O_2_-treated conditions by monitoring OD_600_ values over time. In untreated conditions, all three strains the wild-type *F. tularensis* LVS, the Δ*trxB* mutant, and the transcomplemented strain demonstrated normal growth rates and no growth defects. However, when exposed to H_2_O_2_, the wild-type *F. tularensis* LVS exhibited a slower growth rate than in untreated conditions yet still showed a gradual increase in OD_600_ values over time. In contrast, the Δ*trxB* mutant displayed minimal growth following H_2_O_2_ treatment, suggesting an enhanced susceptibility to oxidative stress relative to the wild-type *F. tularensis* LVS. The transcomplemented strain showed improved growth compared to the Δ*trxB* mutant although the OD_600_ values did not reach the levels observed for the wild-type *F. tularensis* LVS (**Fig. 1A**).

**FIGURE 1:**
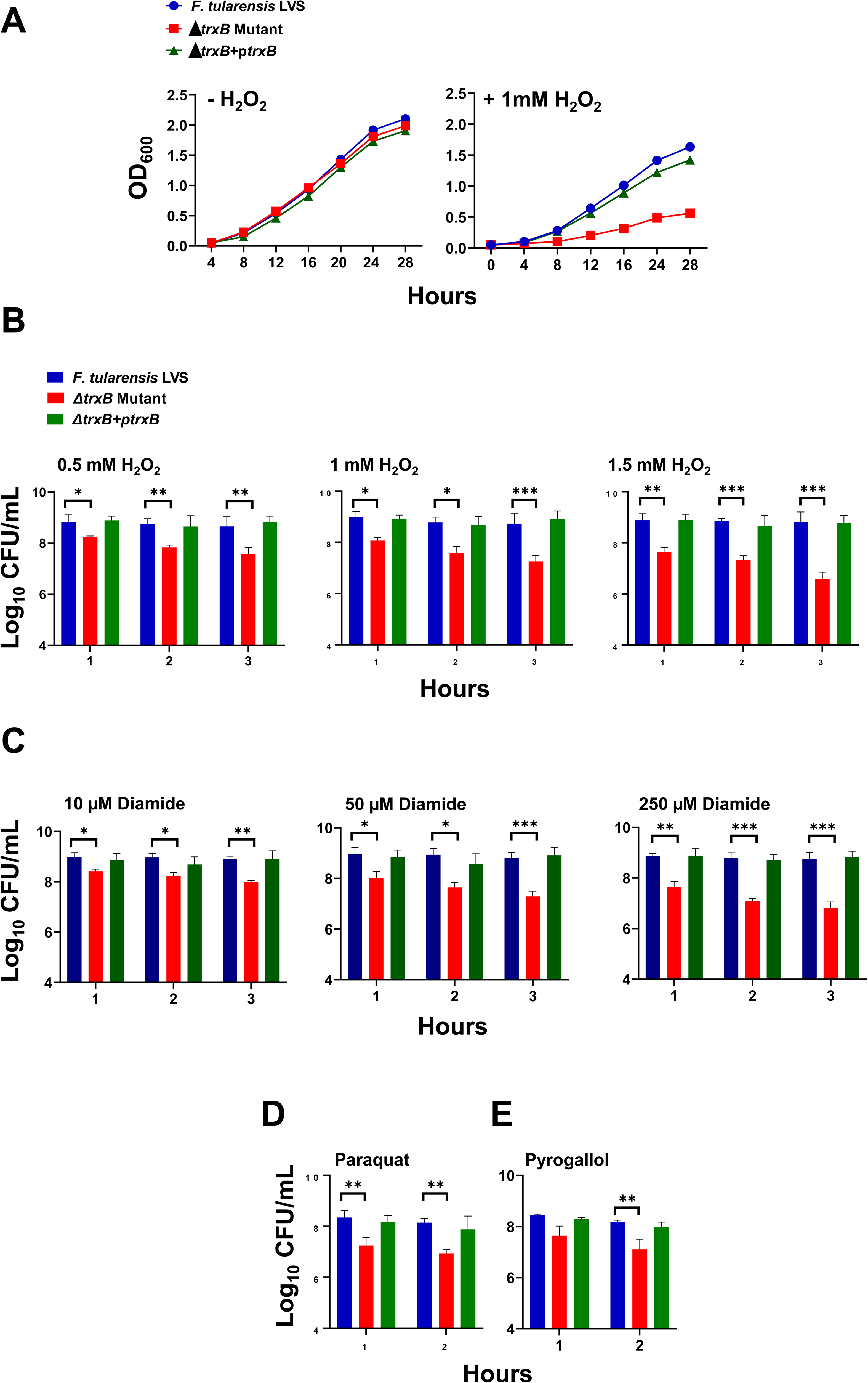
TrxB contributes to the oxidative stress resistance of *F. tularensis*. **(A)** The growth curves of *F. tularensis* LVS, Δ*trxB* mutant, and transcomplemented strain (Δ*trxB*+p*trxB*) in the absence or the presence of 1mM hydrogen peroxide (H_2_O_2_). Bacterial killing assay of the *F. tularensis* LVS, Δ*trxB* mutant, and transcomplemented strain (Δ*trxB*+p*trxB*) in the presence of indicated concentrations of H_2_O_2_ **(B)**, diamide **(C),** paraquat (1mM) **(D),** and pyrogallol (1mM) **(E)** at the indicated times post-exposure. The data shown in A are representative of three independent experiments. The results in B-E are represented as colony forming units (CFU) per mL and are cumulative of three independent experiments, each conducted with three technical replicates, and are shown as Mean±SEM. The data were analyzed using one-way ANOVA. \**p*<0.05, \*\**p*<0.01, \*\*\**p*<0.001.

The sensitivity of wild-type *F. tularensis* LVS, the *ΔtrxB* mutant, and the trans-complemented strain to oxidative stress were further evaluated by exposing the bacteria to 0.5, 1, and 1.5 mM H_2_O_2_, or 10, 50, and 250µM of diamide for 1, 2, and 3 hours, and 1mM paraquat and pyrogallol for 1 and 2 hours. Across all conditions and time points, the *ΔtrxB* mutant exhibited significantly reduced viability as indicated by significantly lower colony forming units (CFUs) compared to the wild-type *F. tularensis* LVS and the transcomplemented strain, indicating its increased susceptibility to oxidative stress caused by these oxidants (**Fig. 1B, C, D** and **E**). These findings demonstrate that the *trxB* gene is required for oxidative stress resistance in *F. tularensis*.

### The Δ*trxB* mutant exhibits enhanced sensitivity to several antibiotics

Antibiotic susceptibility of the Δ*trxB* mutant was compared to the wild-type *F. tularensis* LVS and the transcomplemented strain using a conventional disk diffusion assay. No differences in susceptibility between the Δ*trxB* mutant, the wild-type *F. tularensis* LVS, and the transcomplemented strain were observed for ampicillin, carbenicillin, streptomycin, erythromycin, gentamycin, chloramphenicol, and tetracycline. Against ciprofloxacin and levofloxacin, the Δ*trxB* mutant exhibited slightly increased sensitivity (9.13 ± 0.31 mm and 8.27 ± 0.50 mm, respectively) compared to the wild-type and the transcomplemented strain (both 6 ± 0 mm), suggesting enhanced susceptibility to these antibiotics. For nitrofurantoin, the wild-type and the transcomplemented strains showed similar levels of susceptibility (35.17 ± 1.69 mm and 35.58 ± 1.26 mm, respectively), whereas the Δ*trxB* mutant displayed a reduced zone of inhibition (28.14 ± 3.06 mm), indicating decreased sensitivity. Nalidixic acid exposure revealed a greater zone of inhibition for the Δ*trxB* mutant (39.66 ± 0.88 mm) compared to the wild-type *F. tularensis* LVS (30.38 ± 1.11 mm) and the transcomplemented strain (31.00 ± 1.37 mm), indicating statistically significant enhanced susceptibility in the mutant (**Table 1**). These results suggest that the loss of the *trxB* gene alters susceptibility to specific antibiotics.

**Table 1:** Antibiotic susceptibility of the Δ*trxB* mutant of *F. tularensis* LVS.

| Antibiotic | Concentration<br>($\mu\text{g}$/per disk) | *F. tularensis* LVS | $\Delta trxB$ Mutant | $\Delta trxB + ptrxB$ |
| --- | --- | --- | --- | --- |
| Ampicillin | 2 | $6 \pm 0^*$ | $6 \pm 0$ | $6 \pm 0$ |
| Carbenicillin | 100 | $6 \pm 0$ | $6 \pm 0$ | $6 \pm 0$ |
| Streptomycin | 10 | $25.66 \pm 0.51$ | $25.42 \pm 0.57$ | $25.62 \pm 0.64$ |
| **Ciprofloxacin** | 5 | $6 \pm 0$ | **$9.13 \pm 0.31^{**}$** | $6 \pm 0$ |
| **Levofloxacin** | 5 | $6 \pm 0$ | **$8.27 \pm 0.50$** | $6 \pm 0$ |
| **Nitrofurantoin** | 100 | $35.17 \pm 1.69$ | **$28.14 \pm 3.06$** | $35.58 \pm 1.26$ |
| **Nalidixic acid** | 30 | $30.38 \pm 1.11$ | **$39.66 \pm 0.88$** | $31.00 \pm 1.37$ |
| Chloramphenicol | 30 | $36.83 \pm 1.10$ | $36.41 \pm 0.91$ | $36.69 \pm 1.85$ |
| Tetracycline | 30 | $30.51 \pm 1.41$ | $30.13 \pm 2.06$ | $30.68 \pm 1.12$ |
| Gentamycin | 10 | $31.13 \pm 0.41$ | $31.36 \pm 1.75$ | $31.16 \pm 1.31$ |
| Erythromycin | 15 | $6 \pm 0$ | $6 \pm 0$ | $6 \pm 0$ |

\*Zone of inhibition in millimeters.
\*\*Antibiotics in the bold letters had a significantly larger zone of inhibition than wild-type *F. tularensis* LVS. $P < 0.05$ by one-way ANOVA.

The Δ*trxB* mutant of *F. tularensis* does not exhibit enhanced sensitivity to auranofin.

Auranofin, a gold-containing compound, is known to target specifically the TrxR enzymes in bacteria, inhibiting their activity. Previous studies have demonstrated that bacterial strains lacking TrxR exhibit resistance to auranofin (26, 27). We investigated whether TxrB serves as a target of auranofin in *F. tularensis* by performing growth curve and bacterial killing assays. While exposure to increasing concentrations of auranofin resulted in reduced bacterial growth across wild-type *F. tularensis* LVS, the *ΔtrxB* mutant, and the transcomplemented strain compared to untreated controls, no significant differences in sensitivity were observed among the three strains (**Fig. 2A**). Moreover, identical numbers of viable bacteria were recovered for all three strains at 1-, 3-, and 6-hours following exposure to increasing concentrations of auranofin (**Fig. 2B**). These findings indicate that in contrast to other bacterial species, the TxrB in *F. tularensis* is not a functional target of auranofin.

**FIGURE 2:**
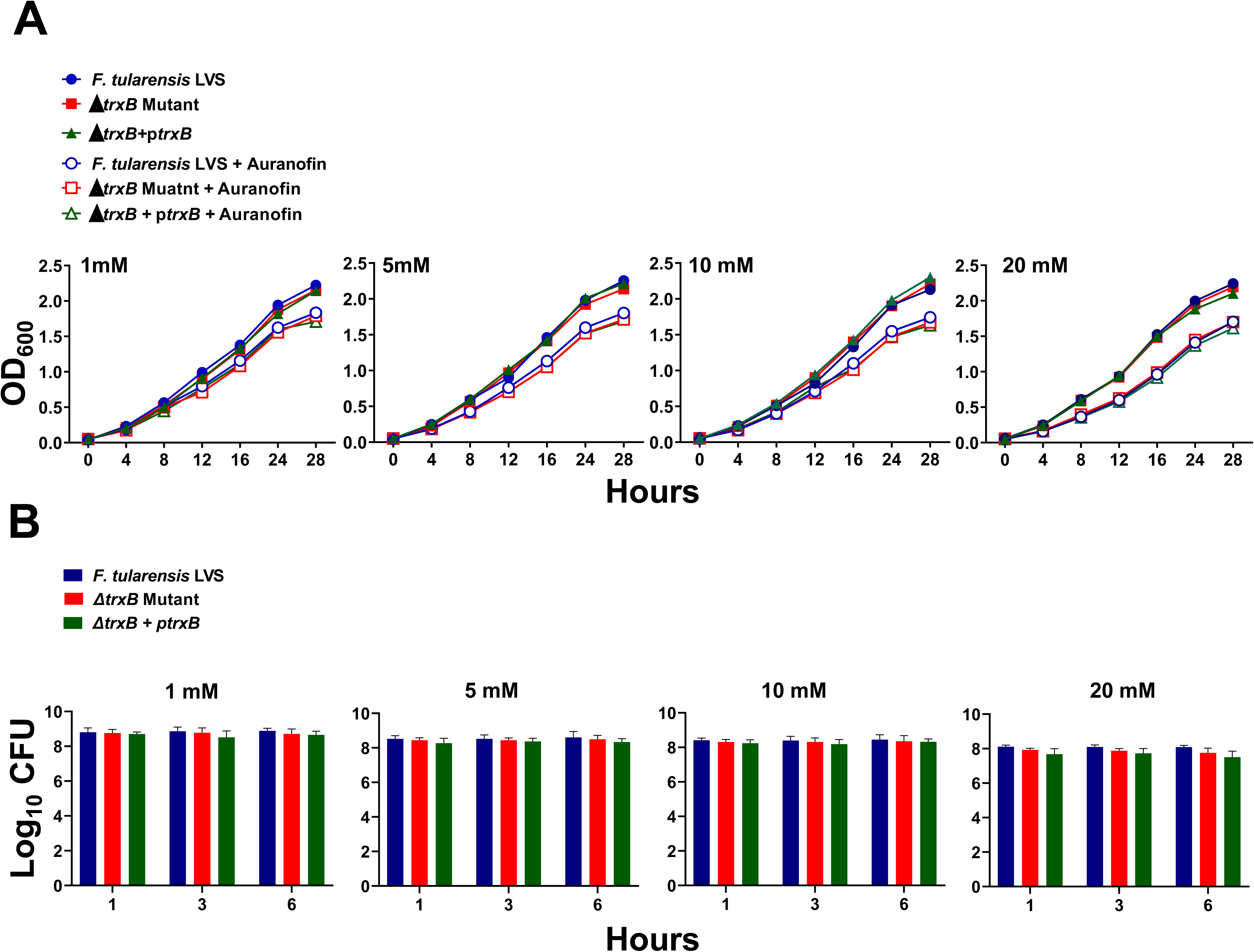
The Δ*trxB* mutant of *F. tularensis* does not exhibit enhanced sensitivity to auranofin. The growth curves **(A)** and bacterial killing assays **(B)** of *F. tularensis* LVS, Δ*trxB* mutant, and transcomplemented strain (Δ*trxB*+p*trxB)* in the absence or the presence of indicated concentrations of auranofin. The data shown in A are representative of three independent experiments. The results in B are represented as colony forming units (CFU) per mL and are cumulative of three independent experiments, each conducted with three technical replicates, and are shown as Mean±SEM. The data were analyzed using one-way ANOVA.

### Expression of oxidative stress response genes and proteins are downregulated in the Δ*txrB* mutant when exposed to oxidative stress caused by H_2_O_2_ and diamide

We next investigated the role of TxrB in the oxidative stress response of *F. tularensis* upon exposure to oxidative stress inducers H_2_O_2_ and diamide. The transcriptional profiles of the oxidative stress response genes alkyl hydroperoxyreductase C (*ahpC*), catalase (*katG*), and superoxide dismutase B (*sodB*), as well as the oxidative stress response regulator gene (*oxyR*), were assessed under both non-stress (-H_2_O_2_) and oxidative stress (+ H_2_O_2_) conditions in the wild type *F. tularensis* LVS, the Δ*trxB* mutant, and the transcomplemented strains (**Fig. 3A**). Under non-stress conditions, the expression levels of all genes remained consistent across the wild-type *F. tularensis* LVS, the Δ*trxB* mutant, and the transcomplemented strains, with no significant differences observed. Following exposure to oxidative stress induced by H_2_O_2_, the expression of primary antioxidant enzyme genes *ahpC, katG, sodB*, and *oxyR* were significantly downregulated in the Δ*trxB* mutant compared to the wild-type *F. tularensis* LVS and the transcomplemented strain (**Fig. 3A**). Western blot analysis corroborated these findings, showing markedly reduced levels of KatG and SodB proteins in the Δ*trxB* mutant relative to the wild-type *F. tularensis* LVS or the transcomplemented strain. FopA was used as a loading control, and demonstrated uniform expression across all conditions (**Fig. 3B**). Identical results were observed in response to the oxidative stress induced by diamide (**Fig. 4A and 4B**). Collectively, these findings highlight the critical role of TrxB in regulating the oxidative stress response in *F. tularensis* LVS and demonstrate that its loss is associated with reduced expression of antioxidant enzyme genes leading to an enhanced sensitivity to oxidative stress.

**FIGURE 3:**
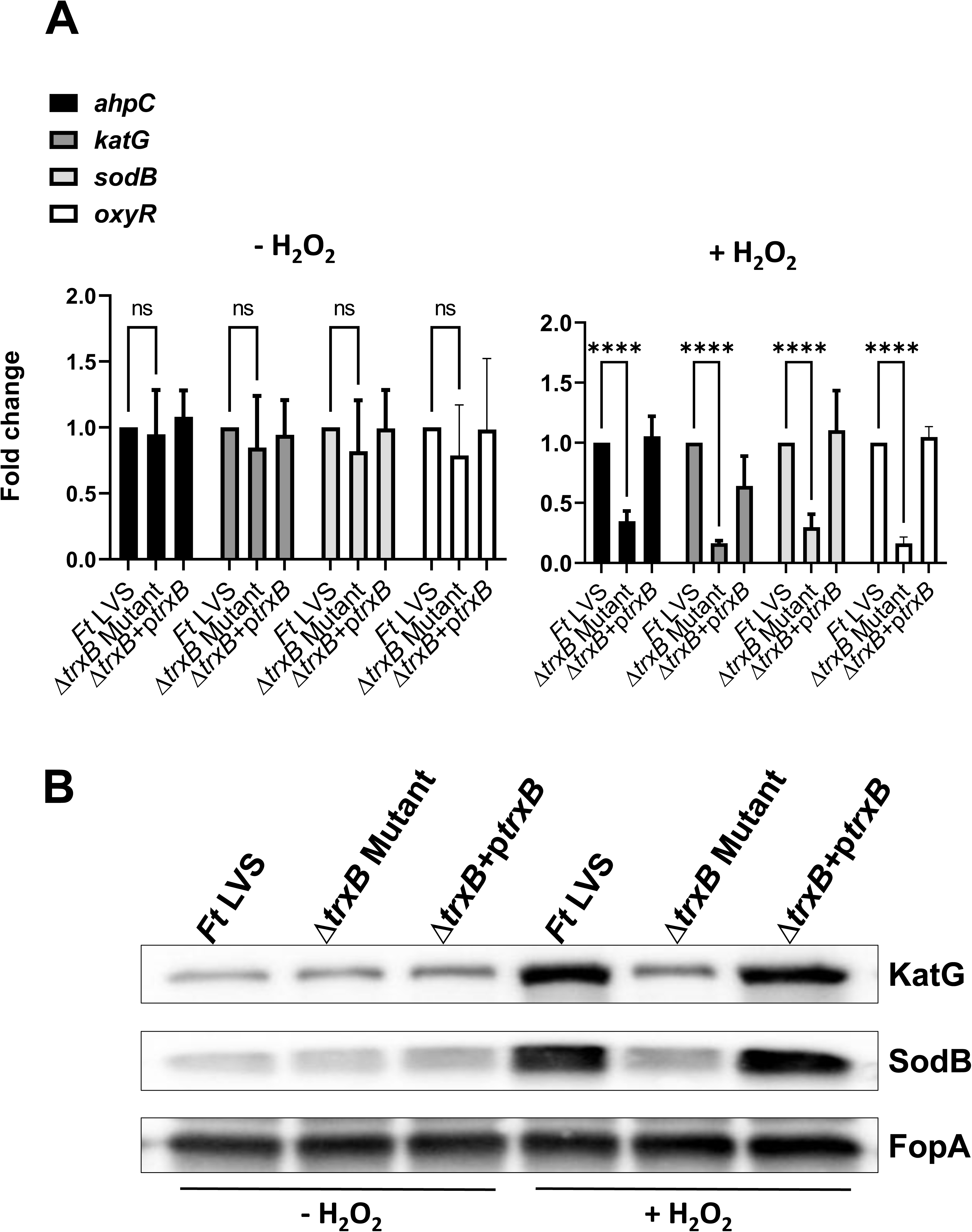
Expression of oxidative stress response genes and proteins are downregulated in the Δ*txrB* mutant when exposed to oxidative stress caused by H_2_O_2_. (A) **Wild-type *F*.** *tularensis* LVS, Δ*trxB* mutant, and the transcomplemented strain (Δ*trxB*+p*trxB*) were either left untreated or exposed to 1 mM H_2_O_2_ for 2 hours to induce oxidative stress. The RNA was isolated and the expression levels of *ahpC*, *katG*, *sodB*, and *oxyR* genes were quantitated by qRT-PCR. The data are represented as the relative fold-change compared to wild-type *F. tularensis* LVS. The data shown are cumulative of three independent experiments, each conducted with three technical replicates, expressed as Mean±SEM, and analyzed using one-way ANOVA. \*\*\*\**p*<0.0001. **(B)** The western blot analysis of the lysates of the indicated *Francisella* strains in the absence or the presence of 1 mM H_2_O_2_ probed with anti-*Francisella* KatG antibodies. The blots were stripped and re-probed with anti-*Francisella* sodB antibodies. The blots were stripped again and re-probed with anti-*Francisella* FopA antibodies that served as loading controls. The results are representative of two independent experiments conducted.

**FIGURE 4:**
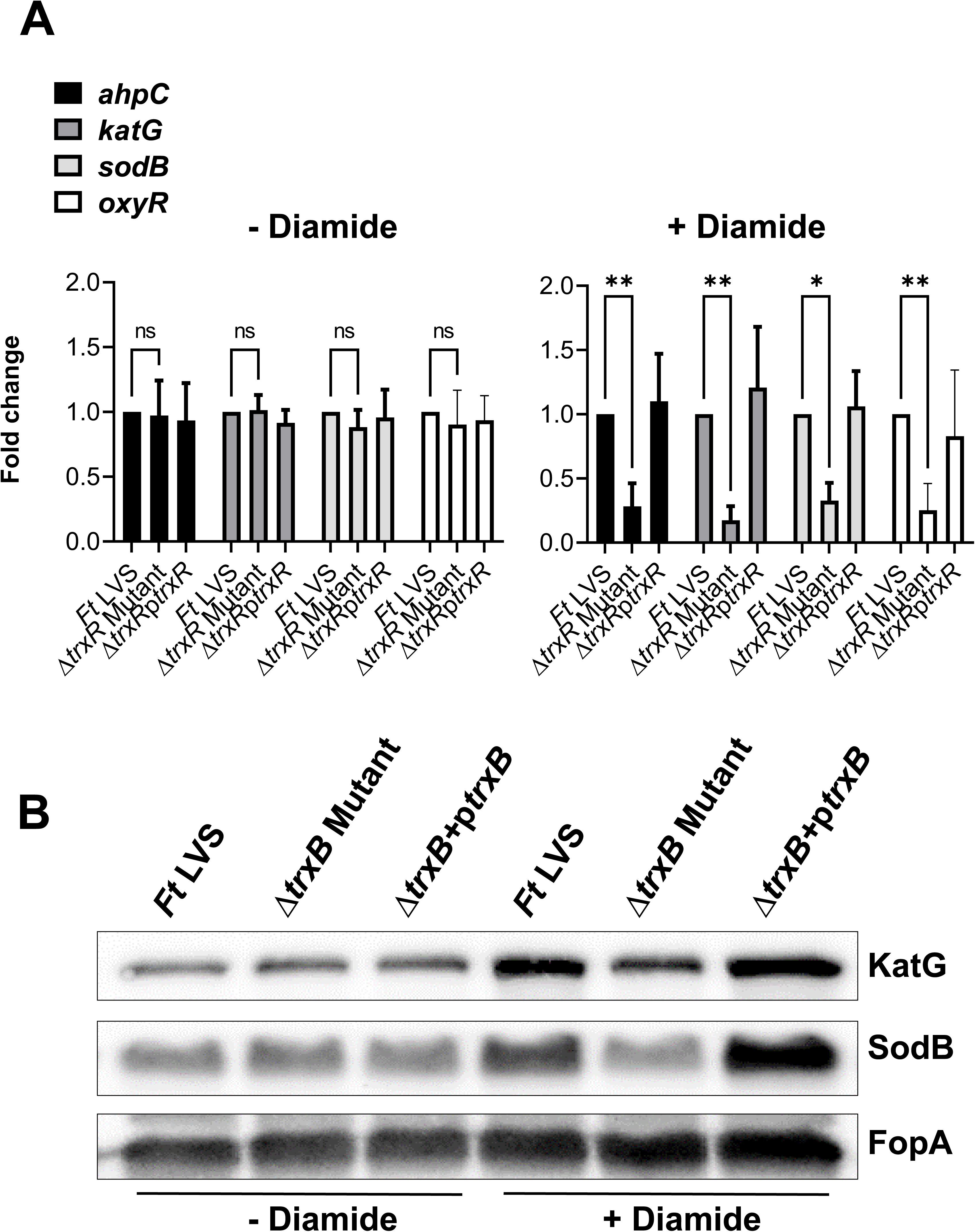
Expression of oxidative stress response genes and proteins are downregulated in the Δ*txrB* mutant when exposed to oxidative stress caused by diamide. **(A)** Wild-type *F. tularensis* LVS, Δ*trxB* mutant, and the transcomplemented strain (Δ*trxB*+p*trxB*) were either left untreated or exposed to 50 µM of diamide for 2 hours to induce oxidative stress. The RNA was isolated and the expression levels of *ahpC*, *katG*, *sodB*, and *oxyR* genes were quantitated by qRT-PCR. The data are represented as the relative fold-change compared to wild-type *F. tularensis* LVS. The data shown are cumulative of three independent experiments, each conducted with three technical replicates, expressed as Mean±SEM, and analyzed using one-way ANOVA. \**p*<0.05, \*\**p*<0.01. **(B)** The western blot analysis of the lysates of the indicated *Francisella* strains in the absence or the presence of 50 µM of diamide probed with anti-*Francisella* KatG antibodies. The blots were stripped and re-probed with anti-*Francisella* sodB antibodies. The blots were stripped again and re-probed with anti-*Francisella* FopA antibodies that served as loading controls. The results are representative of two independent experiments conducted.

### The expression TrxB is regulated by OxyR in *F. tularensis* LVS

Since we observed that the expressions of major OxyR-regulated antioxidant enzyme genes *ahpC, katG,* and *sodB* were downregulated in the Δ*trxB* mutant under the conditions of oxidative stress, we next investigated if the *trxB* gene is regulated by OxyR in *F. tularensis* LVS. We determined the expression profile of the thioredoxin genes *trxA1* and *trxA2*, which are not regulated by OxyR (23), along with the *txrB* gene in wild-type *F. tularensis* LVS, the Δ*oxyR* mutant and the transcomplemented strain. In the absence of oxidative stress, no differences in the expression of *trxA1, trxA2,* and *trxB* genes were observed. However, the expression of the *trxB* gene was significantly downregulated in the Δ*oxyR* mutant, in the presence of oxidative stress induced by H_2_O_2_, while the expression of *trxA1* and *trxA2* genes remained unaltered (**Fig. 5**). These results indicated that the expression of *trxB* is regulated by OxyR.

**FIGURE 5:**
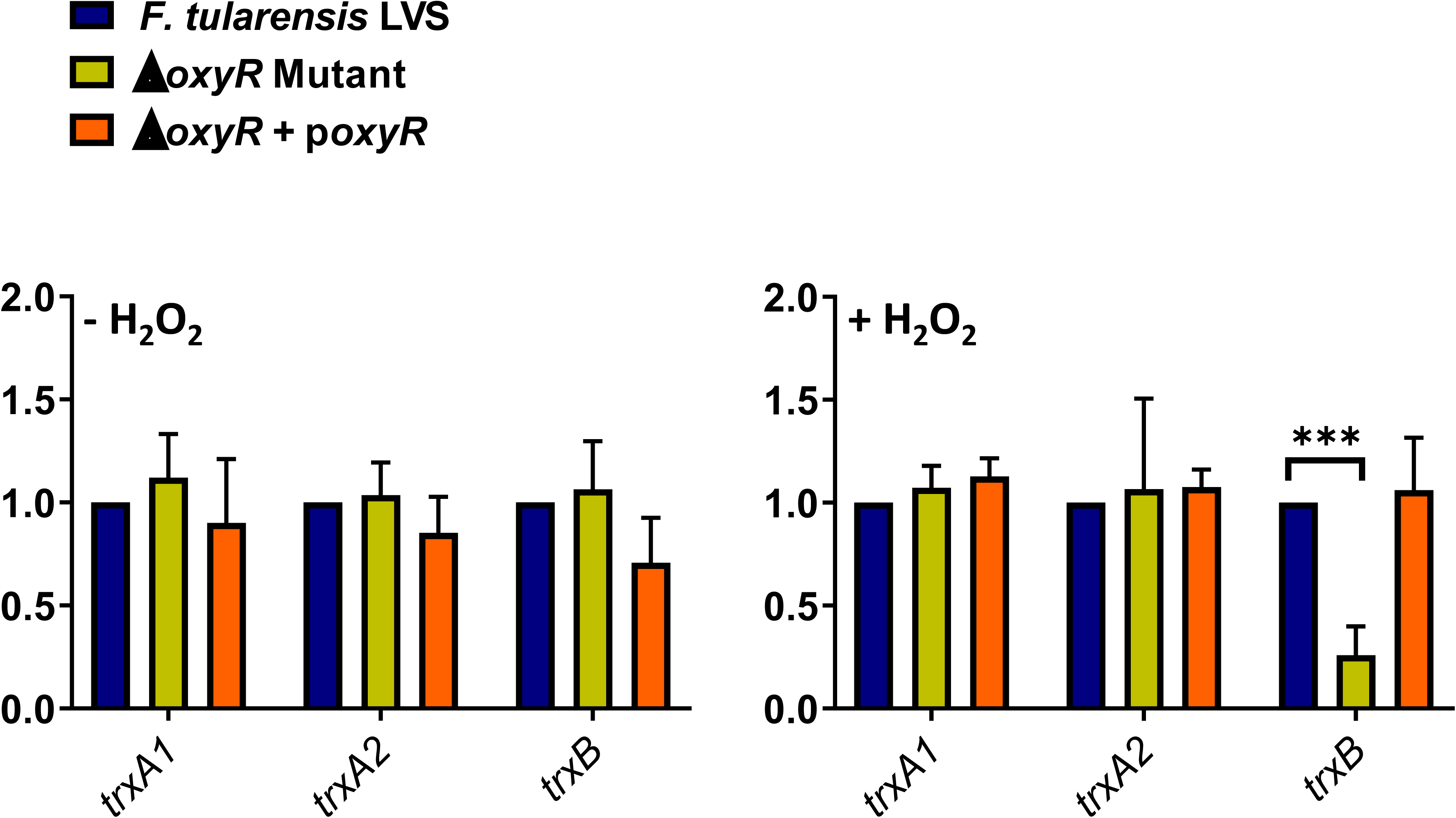
The expression TrxB is regulated by OxyR in *F. tularensis* LVS. Wild-type *F. tularensis* LVS, Δ*oxyR* mutant, and the transcomplemented strain (Δ*oxyR*+p*oxyR*) were either left untreated or exposed to 1 mM H_2_O_2_ for 2 hours to induce oxidative stress. The RNA was isolated and the expression levels of thioredoxin A1 (*trxA1*), thioredoxin A2 (*trxA2*), and thioredoxin reductase (*trxB*) genes were quantitated by qRT-PCR. The data are represented as the relative fold-change compared to wild-type *F. tularensis* LVS. The data shown are cumulative of three independent experiments, each conducted with three technical replicates, expressed as Mean±SEM, and analyzed using one-way ANOVA. \*\*\**p*<0.0001.

We further investigated how OxyR, the master regulator of oxidative stress, regulates the expression of *trxB* in *F. tularensis* LVS. We investigated if OxyR regulates the expression of *trxB* by binding directly to the promoter region of *trxB* by ChIP and EMSA analyses. The *trxB* promoter region was analyzed for critical binding sequences (**Fig. 6A**). Binding sites upstream of the *trxB* gene (F1) and in the intergenic region between the *FTL_1570* and the *trxB* gene (F2) and one site within the *trxB* gene (F3) were tested for OxyR interaction. ChIP assays revealed significant OxyR binding to the *trxB* gene promoter, specifically within the F2 intergenic region (**Fig. 6B**). No binding was observed in the mock controls, confirming the direct interaction of OxyR with the *trxB* promoter. The *F. tularensis* LVS *rpoC*-VSVG strain, which served as an additional control, demonstrated no nonspecific binding in adjacent regions. To validate and further explore OxyR binding specificity, EMSA assays were conducted using the *trxB* gene promoter probes. In the wild-type *F. tularensis* LVS, OxyR bound specifically to the *trxB* promoter probe, an interaction that was disrupted upon the addition of competitor DNA, confirming its sequence specificity. Binding was notably absent in the Δ*oxyR* mutant strain, whereas the Δ*oxyR*+p*oxyR* transcomplemented strain restored the OxyR-*trxB* promoter binding (**Fig. 6C**). As an additional control, EMSA analysis using PmrA protein and its cognate *pmrA* promoter was conducted. PmrA bound exclusively to its own promoter and exhibited no interaction with the *trxB* promoter probes, demonstrating the specificity of the OxyR-*trxB* promoter interaction (**Fig. 6D**). Together, these findings demonstrate that OxyR directly binds to the *trxB* promoter. The specificity of OxyR-*trxB* promoter interaction and its absence in the Δ*oxyR* mutant further establish the role of OxyR as a key transcriptional regulator of *trxB* in *F. tularensis* LVS.

**FIGURE 6:**
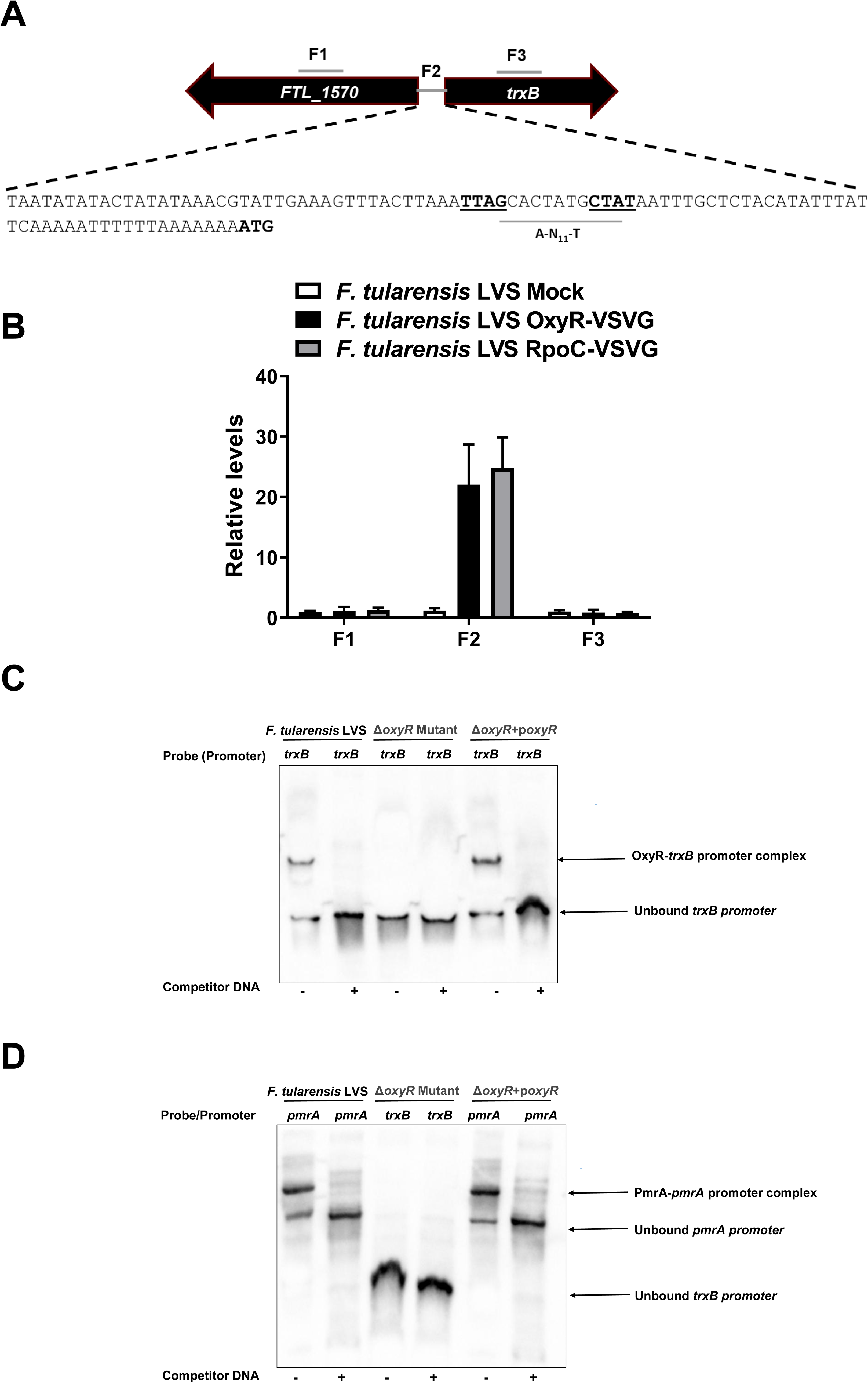
OxyR in *F. tularensis* LVS regulates the expression of TrxB. (A, **B)** Chromatin immunoprecipitation (ChIP) was performed using anti-VSV-G-agarose beads. The fragments of intergenic regions covering the putative promoter sequences and the coding regions of the genes up and downstream of the *trxB* gene (indicated by numbers) in ChIP and input samples of the indicated bacterial strains were analyzed for enrichment by qRT-PCR. The values were normalized to the input and a *fopA* coding region as internal controls. The qRT-PCR results of ChIP from *F. tularensis* LVS *oxyR-VSVG* along with wild-type *F. tularensis* LVS (mock) and *rpoC-VSVG* (positive control) are shown (n=3 biological replicates). The data shown are representative of three independent experiments and were analyzed by ANOVA. **(C)** Electrophoretic mobility shift assay (EMSA) with the promoter region for *trxB* of *F. tularensis.* EMSA was performed using bacterial lysates from the *F. tularensis* LVS, the Δ*oxyR* mutant, and the transcomplemented strain (Δ*oxyR*+p*oxyR*). The biotinylated DNA sequence of the promoter region was used as the probe, whereas the unlabeled promoter region was used as competitor DNA. **(D)** EMSA with the promoter region for the *pmrA* gene of *F. tularensis.* The activity of lysates from the Δ*oxyR* mutant was determined by binding of transcriptional regulator PmrA to its putative *pmrA* promoter region by EMSA. A biotinylated 505 bp fragment of the *pmrA* promoter region was used as a probe, whereas an unlabeled promoter region was used as competitor DNA.

### The Δ*trxB* mutant is attenuated for intramacrophage survival and its clearance is dependent on ROS

We performed a macrophage invasion assay in murine macrophage cell line RAW264.7 to determine the role of TrxB in macrophage survival. Almost equal numbers of wild-type *F. tularensis* LVS, the Δ*trxB* mutant, and the transcomplemented bacteria were recovered from the infected macrophages after 4 hours of infection. However, nearly 100-fold less Δ*trxB* mutant as compared to the wild-type *F. tularensis* LVS and the transcomplemented bacteria recovered at 24 hours post-infection (**Fig. 7A**). In BMDMs derived from the wild-type C57BL/6 mice, the numbers of Δ*trxB* mutant bacteria recovered at 4- and 24 hours post-infection were significantly lower than those observed for the wild-type *F. tularensis* LVS or the transcomplemented strain. However, the Δ*trxB* mutant bacteria replicated similarly to the wild-type *F. tularensis* LVS and the transcomplemented strain when the assays were conducted using *gp91phox^-/-^* BMDMs that cannot generate the ROS (**Fig. 7B**). Collectively, these results demonstrate that TrxB contributes to the intramacrophage survival of *Francisella* by overcoming ROS-induced oxidative stress.

**FIGURE 7:**
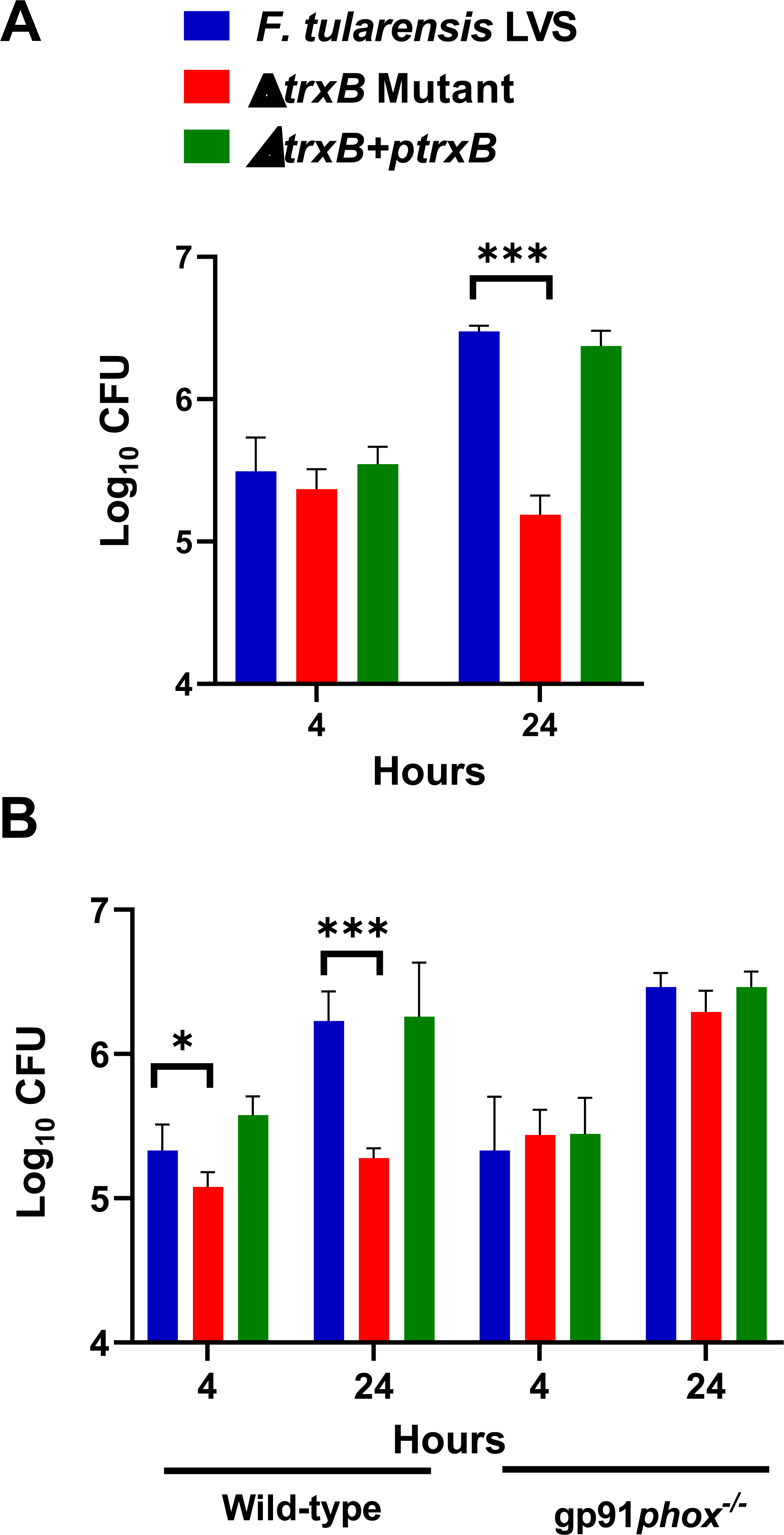
The Δ*trxB* mutant is attenuated for intramacrophage survival and its clearance is dependent on reactive oxygen species. Murine macrophage cell line RAW264.7. **(A)** or primary BMDMs isolated from wild-type C57BL/6 mice or gp91*Phox*^−/−^ (B) were infected with the *F. tularensis* LVS, the Δ*trxB* mutant or the transcomplemented strain (Δ*trxB*+p*trxB*) at 100 MOI of infection. The intramacrophage growth was quantitated at 4- and 24-hour post-infection. The results are expressed as Log_10_ CFU/mL. The data represented as Mean±SEM are cumulative of three independent experiments, each conducted with four technical replicates and analyzed using one-way ANOVA. *** *P*<0.001.

## Discussion

Thioredoxin reductases (TrxRs) are classified into high molecular weight (HMW) TrxRs and low molecular weight (LMW) TrxRs based on their molecular weights (28). In addition to the molecular weights, these TrxRs differ in their structural complexity and subunit organization. HMW TrxRs are dimeric enzymes comprising two subunits, predominantly found in mammals as selenium-containing proteins requiring selenium for their catalytic activity. In contrast, LMW TrxRs are predominantly monomeric and exhibit simpler structures. They lack selenium and are widely distributed among bacteria, plants, and fungi (29, 30). Both types of TrxRs catalyze the reduction of oxidized thioredoxin to its reduced form, a key reaction in maintaining cellular redox homeostasis. However, despite their functional similarities, HMW and LMW TrxRs may exhibit distinct substrate specificities and affinities. HMW TrxRs are required for various cellular processes, including DNA synthesis and repair. The LMW TrxRs mediate oxidative stress responses and maintain cellular redox homeostasis in bacteria, regulate the enzymatic activity of the thiol-containing enzymes by maintaining those in their functional reduced state, stabilize protein structures and functions by forming and repairing disulfide bonds, and in signal transduction by modulating the redox state of the signaling proteins.

TrxRs are NADPH-dependent enzymes that maintain redox homeostasis and contribute to bacterial virulence. These enzymes are essential for bacterial adaptation to oxidative stress, such as ROS generated during aerobic growth or by immune responses (31). In *M. tuberculosis*, the thioredoxin reductase, TrxB2, is required to resist thiol-oxidizing stress and establish and maintain infection in mice (32). In *H. pylori*, they regulate the redox state of proteins that are necessary for colonization and immune evasion (33). In *E. coli* and *S. aureus*, TrxRs regulate enzymes essential for bacterial survival under oxidative stress (34, 35). In *Clostridium difficile* and *Listeria monocytogenes*, TrxRs are involved in regulating virulence and repairing oxidative damage (36, 37). In this study, we demonstrate that the TrxR denoted as TrxB of *F. tularensis* LVS is critical for resistance to oxidative stress, antibiotics, and intramacrophage survival. Additionally, it reveals the complex antioxidative stress resistance mechanisms in *F. tularensis* and how their regulation systems are interconnected.

The amino acid sequence alignment revealed that TrxRs exhibit conserved functional motifs that are essential for their enzymatic activity and are consistently found across various bacterial pathogens, including *F. tularensis*. The most notable motif is the active site containing the conserved sequence CXXC, where the cysteine residues facilitate thiol-disulfide exchange reactions critical for reducing oxidized thioredoxin. Another key motif is the flavin adenine dinucleotide (FAD)-binding domain, which supports the transfer of electrons from NADPH to the active site cysteines (22). The presence of these conserved features across diverse bacterial species highlights their role in maintaining cellular redox balance and adapting to oxidative stress.

The results from this study demonstrate that the Δ*trxB* mutant of *F. tularensis* LVS is sensitive to oxidative stress induced by H₂O₂ and superoxide-generating compounds, paraquat and pyrogallol. These oxidants generate ROS that oxidize thiol groups in proteins, leading to the formation of disulfide bonds and disruption of cellular functions. Our previous studies have shown that *F. tularensis* encodes two thioredoxins, TrxA1 and TrxA2. However, the bacterium primarily relies on TrxA1 to counter oxidative stress and ensure intracellular survival (23). For H₂O₂, thioredoxins work in conjunction with peroxiredoxins such as AhpC to detoxify the oxidant, converting it into water and protecting cellular components from oxidative damage. TrxRs then catalyze the NADPH-dependent reduction of oxidized thioredoxins, restoring their ability to reduce disulfide bonds in proteins and repair oxidative damage (38). In the case of paraquat, which generates superoxide radicals, TrxRs indirectly mitigate oxidative stress by maintaining the reduced state of thioredoxin, thereby supporting the activity of antioxidant enzymes like superoxide dismutases (39). Pyrogallol, another ROS generator, induces oxidative stress by producing hydroxyl radicals. Similarly, the oxidative stress induced by diamide, a thiol-specific oxidizing agent, is countered by TrxRs through their role in maintaining the reduced state of thioredoxin (23). Diamide promotes the formation of disulfide bonds in proteins, leading to the oxidation of thiol groups and disruption of cellular redox homeostasis. In bacterial pathogens, TrxRs are upregulated in response to diamide-induced stress, ensuring efficient detoxification of reactive disulfides and supporting survival under oxidative stress conditions (40–42). Thus, TrxRs counter oxidative stress induced by these oxidants by maintaining the reduced state of thioredoxin, which is essential for cellular redox increased sensitivity to these oxidants, as has been observed in this study.

The functional contributions of TrxRs to antibiotic resistance are not uniform across bacterial species and may vary depending on the antibiotic in question and the specific bacterial strain (43). While TrxRs themselves do not directly confer antibiotic resistance, their role in regulating the redox state of enzymes involved in antibiotic metabolism or target binding has been implicated in resistance mechanisms in certain bacterial pathogens. TrxRs have been associated with metronidazole resistance in *H. pylori*, as they modulate the activity of redox-sensitive enzymes required for the bioactivation of metronidazole (44). In *M. tuberculosis*, TrxRs have been linked to resistance against streptomycin, potentially by influencing redox-sensitive pathways critical for drug efficacy (45). Furthermore, TrxRs have been shown to play a role in rifampicin resistance in *S. aureus*, possibly by modulating redox-dependent mechanisms affecting drug binding or metabolism (46). TrxRs also play a role in antibiotic resistance by influencing redox-sensitive enzymes involved in antibiotic detoxification or binding, as seen in species such as *Pseudomonas aeruginosa* (47). Our results demonstrate that the loss of the *trxB* gene alters susceptibility to specific antibiotics, potentially by impacting the redox status of the bacteria. While certain antibiotics, such as ciprofloxacin, levofloxacin, and nalidixic acid, showed significant differences, sensitivity to other antibiotics remained unaffected, emphasizing the selective role of *trxB* in mediating antibiotic resistance of *F. tularensis*.

The sensitivity of several bacteria to a gold-containing compound, auranofin, has been particularly of interest in the context of TrxRs. It has been reported that auranofin specifically targets TrxR in bacterial cells and inhibits its activity. The strains lacking TrxR are resistant to it, while the wild-type bacteria still retain their sensitivity. Auranofin has antibacterial activity against several bacterial species, including *Staphylococcus aureus*, *Streptococcus pneumoniae*, and *Mycobacterium tuberculosis* (48, 49). Initially identified as a TrxR inhibitor due to its ability to disrupt cellular redox balance and induce oxidative stress, auranofin binds to the active site of TrxR, inhibiting its enzymatic activity. However, our results do not conform to these findings and demonstrate that there were no differences in the sensitivity of the wild-type *F. tularensis* LVS or the Δ*trxB* mutant to auranofin used at various concentrations. These findings align with several other findings demonstrating that bacterial TrxRs are not the targets of auranofin. In fact, resistance to auranofin in several Gram-negative bacteria is attributed to outer membrane proteins acting as a barrier or the efficient efflux of the drug via efflux pumps. It has also been suggested that resistance to auranofin could result from an efficient glutathione system (50). *Francisella* possesses all three structural features, and therefore, the lack of antibacterial effects of auranofin may result from any or all of these structural components.

Our results demonstrate that the expression of primary antioxidant genes *ahpC, KatG,* and *sodB* was significantly downregulated in the Δ*trxB* mutant when exposed to oxidative stress induced by H_2_O_2_ or diamide. We have reported that the expression of these primary antioxidant genes is controlled by OxyR, which is the key regulator of oxidative stress response in *F. tularensis* (16). OxyR is activated through the formation of a disulfide bond between conserved cysteine residues. Once activated, OxyR binds to the promoter regions of target genes and induces their transcription. This upregulation enhances the cellular capacity to counteract oxidative damage by maintaining redox homeostasis. In our previous study, we reported that the expression of both thioredoxins TrxA1 and TrxA2 is not regulated by OxyR (23). However, the result from this study demonstrates that the expression of TrxB is directly under the control of OxyR under the conditions of oxidative stress.

To conclude, this study describes the role of an important uncharacterized component of the oxidative stress response machinery of *F. tularensis* required for resistance to oxidative stress and intramacrophage survival.

## Materials and Methods

### Bacterial strains and culture media

The bacterial strains, plasmids, and mutants generated in this study are listed in **Table 2**. The *Francisella tularensis* subsp. *holarctica* live vaccine strain (LVS) obtained from the American Type Culture Collection, Rockville, MD (ATCC 29684) was used in this study. The deletion mutant of the master regulator of the oxidative stress response (Δo*xyR*) and its transcomplemented strain (Δo*xyR*+p*oxyR*) has been previously generated and characterized in our laboratory (16). The *F*. *tularensis* cultures were grown on Mueller-Hinton (MH) chocolate agar plates (BD Biosciences, San Jose, CA) supplemented with IsoVitaleX at 37°C with 5% CO_2_; MH broth (BD Biosciences, San Jose, CA) supplemented with ferric pyrophosphate and IsoVitaleX (BD Biosciences, San Jose, CA) at 37°C with shaking (160 rpm). Active mid-log phase bacteria grown in MH-broth were harvested and stored at −80°C. One mL aliquots were thawed periodically for use. *Escherichia coli* strain DH5α was used for routine cloning. *E. coli* cultures were grown in Luria-Bertani (LB) broth or on LB agar plates. When necessary, kanamycin (25μg/ml), or hygromycin (200 μg/ml) was included in broth and agar media for selection purposes.

**Table 2:**
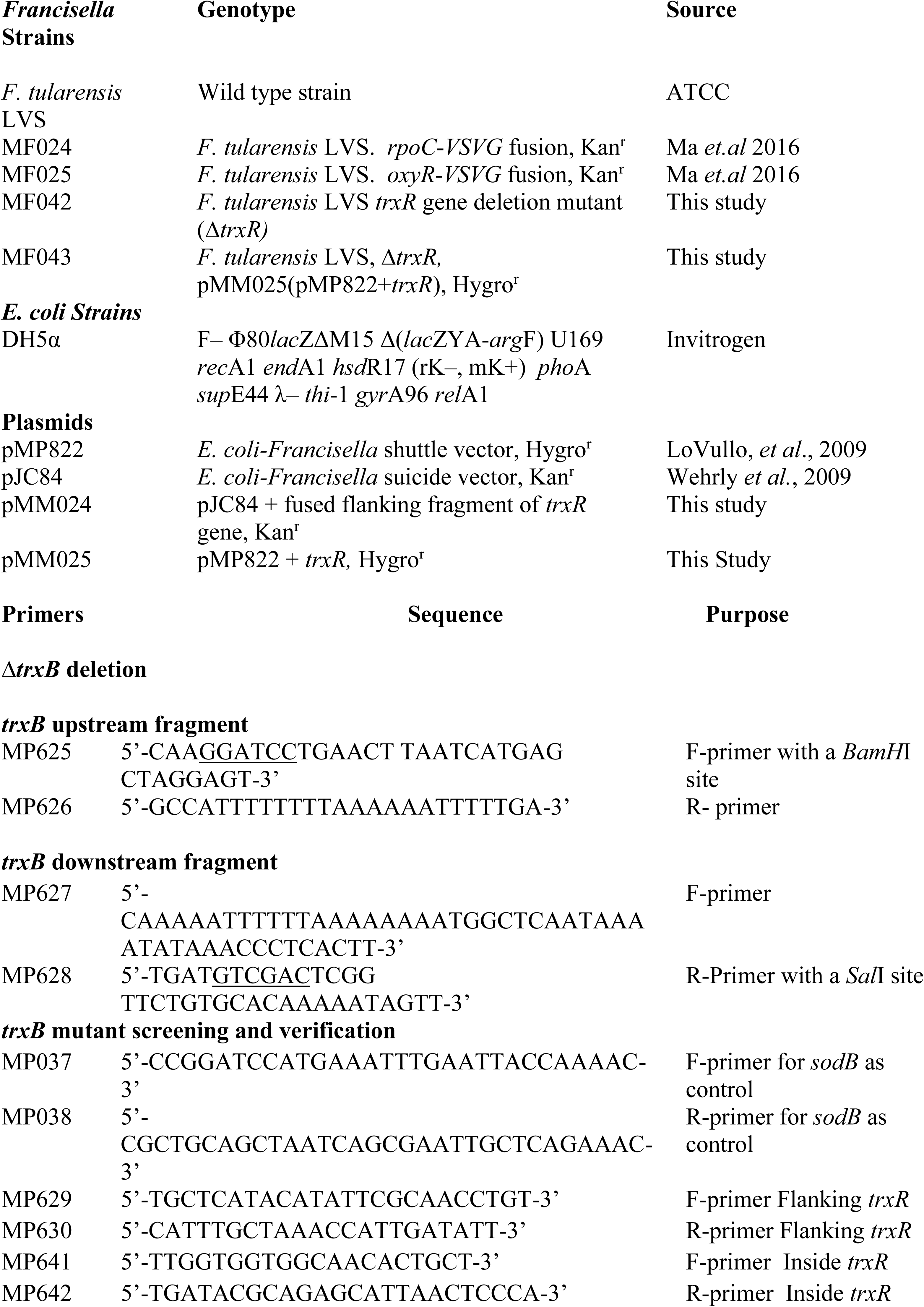
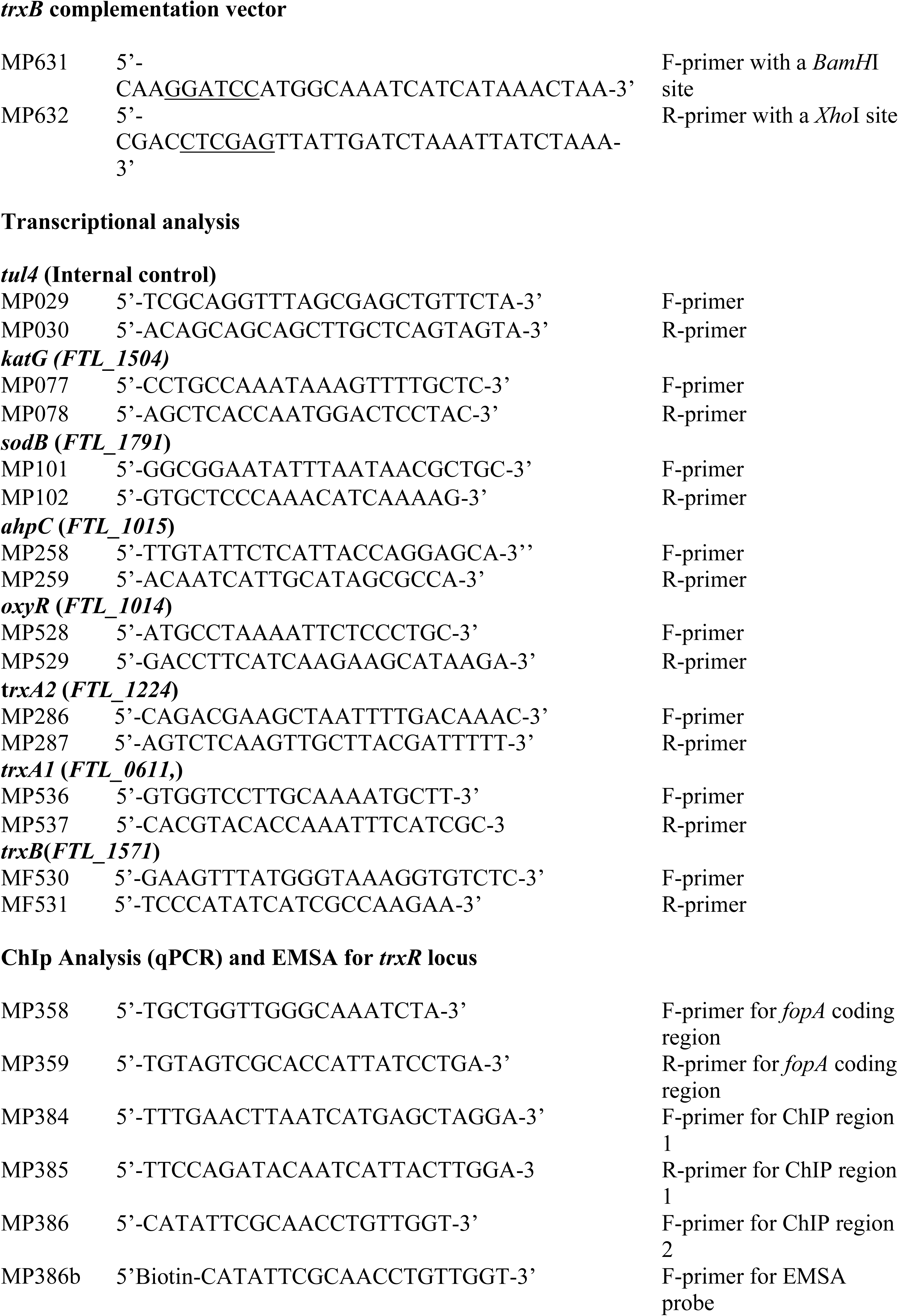
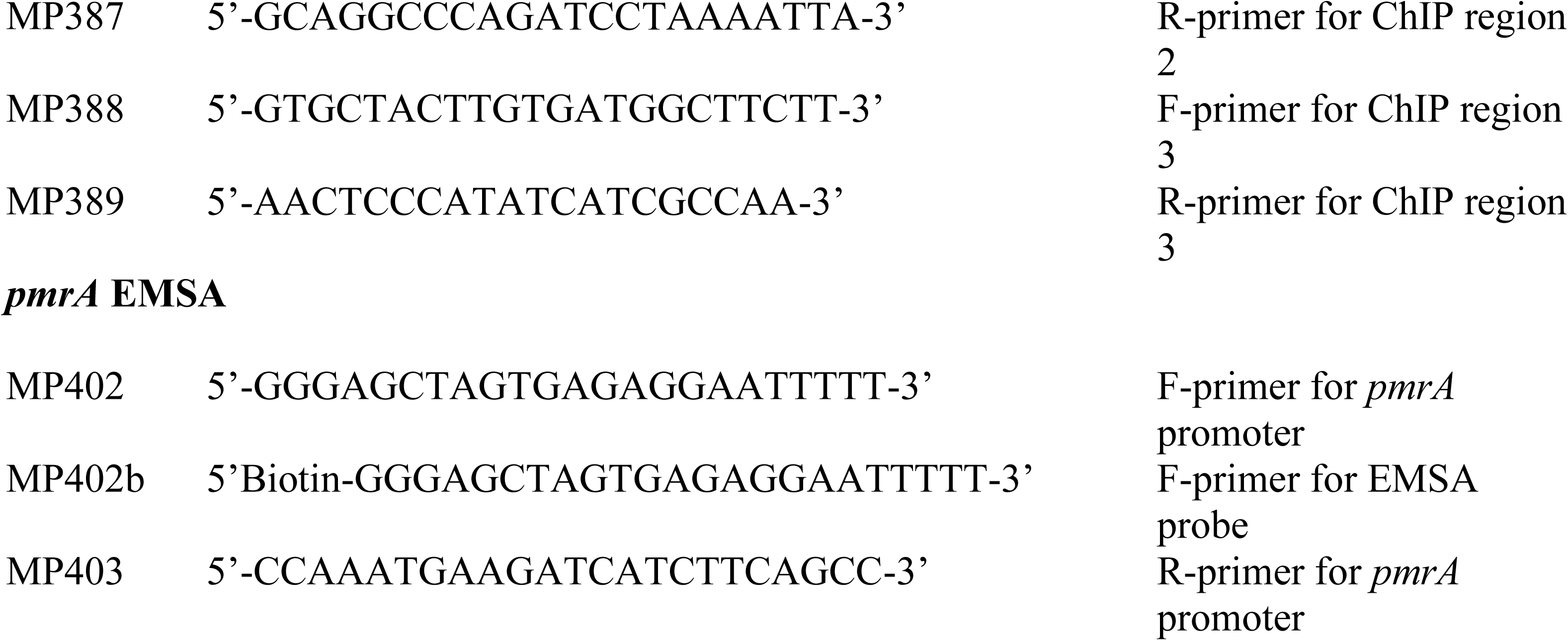
List of bacterial strains, plasmid vectors, and primers used in this study.

### Bioinformatics analysis

Sequences of thioredoxin reductase (TrxR) from *F. tularensis* LVS and other bacterial species were obtained from the National Center for Biotechnology Information (NCBI). Using Clustal Omega, multiple sequence alignments were performed to determine both conserved and diverse features of TrxR across these bacterial species. TrxRs from three subspecies *F. tularensis* subspecies *holarctica* LVS (*FTL_1571, trxB*), *F. tularensis* subspecies *tularensis* SchuS4 (*FTT_0489c*, *trxB*), and *F. novicida* (FTN_0580*, trxB*) were aligned. TrxRs from *E. coli* (*b0888, trxB*), *Salmonella* Typhi (*t1976, trxB*), *Yersinia pestis* (*YPO1374, trxB*), *Helicobacter pylori* (*HP_0825, trxB*), *Pseudomonas aeruginosa* (*PA2616, trxB1*), *Staphylococcus aureus* (*SACOL0829, trxB*), *Bacillus subtilis* (*BSU_34790, trxB*), *Mycobacterium tuberculosis* (*Rv3913, trxB1/trxR*), and *Vibrio cholerae* (*Vch1786_I0686, trxB*) were also included in the analysis.

### Construction of *F. tularensis* LVS *trxB* (*FTL_1571*) gene deletion mutant (Δ*trxB*) and transcomplemented (Δ*trxB* + p*trxB*) strains

An in-frame gene deletion mutant of the *trxB* gene (Δ*trxB*) in *F. tularensis* LVS was constructed using the SacB-assisted allelic replacement vector p*JC84* (51). To create the Δ*trxB* mutant, 850 base pairs (bp) upstream and 954 bp downstream fragments flanking the *trxB* gene were amplified by PCR using primers MP625/626a and MP627/628, respectively, and fused by overlap extension PCR. The resulting 1790 bp fragment was cloned into pJC84 at *BamH*I and *Sal*I sites, generating a plasmid p*MM024*. *F. tularensis* LVS was transformed with p*MM024* by electroporation, as described earlier (16). Following transformation, kanamycin-resistant clones were selected on MH-chocolate agar plates containing 10 µg/ml kanamycin. The kanamycin-resistant clones were plated on MH-chocolate agar plates containing 8% sucrose and the kanamycin-sensitive clones were selected. These clones were screened for allelic replacement by PCR with *trxB* gene-specific primers and confirmed through a duplex PCR using *sodB* gene primers as internal controls, followed by DNA sequencing. To complement the *F. tularensis ΔtrxB* mutant, the *trxB* gene sequence (*FTL_1571*) was PCR amplified using primers MP631 and MP632. The amplified fragment was then digested with *BamH*I and *Xho*I restriction enzymes and cloned into the *E. coli-Francisella* shuttle vector p*MP822* (52). The resulting plasmid, p*MM025*, was verified through PCR and DNA sequencing. Subsequently, p*MM025* was transformed into the *ΔtrxB* mutant by electroporation, and transformants were selected on MH-chocolate agar plates containing 200 µg/mL hygromycin. PCR analysis confirmed the successful complementation of the mutant strain.

### Testing the susceptibility of *ΔtrxB* mutant of *F. tularensis* LVS to oxidants

The wild-type *F. tularensis* LVS, the *ΔtrxB* mutant, and the transcomplemented strains were tested for susceptibility to hydrogen peroxide (H_2_O_2_), diamide, paraquat and pyrogallol using growth curves and bacterial killing assays. For growth curves, bacterial suspensions (OD_600_ = 0.05) were incubated in MH-broth with or without 1 mM H_2_O_2_ at 37°C with constant shaking, and OD_600_ values were recorded every four hours for 28 hours. In bacterial killing assays, bacterial suspensions (1×10⁹ CFU/mL) were exposed to 0.5 mM, 1 mM, and 1.5 mM of H_2_O_2_ or 10, 50 and 250 µM of diamide at 37°C. Similar killing assays were conducted using superoxide-generating compounds paraquat (1 mM), and pyrogallol (1 mM). Samples were taken at 1, 2, and 3 hours, serially diluted, and plated on MH-chocolate agar plates. The plates were incubated at 37°C in the presence of 5% CO_2_ and the colonies were counted 48 hours later. The bacterial counts were expressed as Log_10_ CFU/mL.

### Antibiotic sensitivity assays

The sensitivity of wild-type *Francisella tularensis* LVS, the Δ*trxB* mutant, or the transcomplemented strain to several antibiotics was determined using disc diffusion assays. The bacterial cultures grown on MH chocolate agar plates were scraped and suspended in MH-broth to achieve an OD_600_ of 1.0. The bacterial suspensions were spread with a sterile cotton swab onto MH chocolate agar plates. Sterile antibiotic discs were placed on the plates. The plates were incubated at 37°C in the presence of 5% CO_2_. After 3 days of incubation, the zones of growth inhibition around the disks were measured. The experiments were repeated at least three times for each antibiotic and the bacterial strain.

Both growth curves and killing assays were also performed to test the sensitivity of the wild-type *F. tularensis* LVS, the Δ*trxB* mutant, or the transcomplemented strain to auranofin by growth curves and bacterial killing assays. For growth curves, bacterial suspensions (OD_600_ = 0.05) were incubated in MH-broth with or without 1, 5, 10, and 20 mM auranofin at 37°C with constant shaking, and OD_600_ values were recorded every four hours for 28 hours. In bacterial killing assays, bacterial suspensions (1×10⁹ CFU/mL) were exposed to similar concentrations of auranofin at 37°C. Samples were taken at 1, 3, and 6 hours, serially diluted, and plated on MH-chocolate agar plates. The plates were incubated at 37°C in the presence of 5% CO_2_ and the colonies were counted 48 hours later. The bacterial counts were expressed as Log_10_ CFU/mL.

### Analysis of gene expression

Overnight cultures of wild-type *F. tularensis* LVS, the *ΔtrxB* mutant, and the transcomplemented strain were adjusted to an OD_600_ of 0.2 and grown for 2 hours at 37°C with shaking in 10 mL of MH-broth, both in the absence or presence of 1 mM H_2_O_2_ or 50 µM diamide. Similarly, in another experiment, wild-type *F. tularensis* LVS, the Δ*oxyR* mutant and Δ*oxyR*+p*oxyR* transcomplemented strains were either left untreated or treated with 1mM H_2_O_2_. Total RNA was extracted from the grown cultures using Purelink RNA Mini Kits (Ambion) and further treated with DNase to eliminate contaminating DNA. cDNA synthesis was performed using the iScript cDNA Synthesis Kit (Ambion). Quantitative PCR (qPCR) was then conducted with iQ SYBR Green Supermix (BioRad) to analyze the expression of target genes, using the *tul4* gene of *F. tularensis* LVS as an internal control. Relative gene expression was calculated using the 2−ΔΔCT method: 2− [ΔCT (mutant) − ΔCT (WT)], where ΔCT = CT of the target gene − CT of the internal control. Data were presented as the mean ± SD of three biological replicates. The primer sequences of the target genes used for qPCR are detailed in **Table 2**.

### Western blot analysis

Overnight cultures of wild-type *F. tularensis* LVS, the *ΔtrxB* mutant, and the transcomplemented strain were adjusted to an OD_600_ of 0.2 and grown for 2 hours at 37°C with shaking in 10 mL MH broth in the absence or presence of 1 mM H_2_O_2_ or 50 µM diamide. Bacterial cultures were harvested, centrifuged, and re-suspended in 200 µL of lysis buffer containing 200 mM Tris-HCl (pH 8.0), 320 mM (NH_4_)_2_SO_4_, 5 mM MgCl_2_, 10 mM EDTA, 10 mM EGTA, 20% glycerol, 1 mM dithiothreitol (DTT), and protease and phosphatase inhibitors. Protein concentrations of cell lysates were determined using BioRad reagents. Five micrograms of protein from each sample were separated on a 12% SDS-PAGE gel, transferred to polyvinylidene difluoride (PVDF) membranes (Millipore), and probed with primary antibodies against KatG (1:20,000) and SodB (1:20,000). Secondary anti-rabbit antibodies conjugated to horseradish peroxidase (HRP) (Santa Cruz; 1:10,000) were used for detection. Protein bands were visualized using Supersignal West Pico chemiluminescent substrate (Thermo Scientific) and imaged on a Chemidoc XRS system (BioRad). For loading control, the membranes were stripped and re-probed with anti-FopA antibodies. The anti-SodB, KatG, and FopA antibodies were kindly provided by Dr. Karsten Hazlett (Albany Medical College, Albany, NY).

### Chromatin immunoprecipitation (ChIp) assays

To study the binding activity of OxyR in vivo to the promoter region of the *trxB* gene, an *F. tularensis* LVS *oxyR*-Vesicular Stomatitis Virus Glycoprotein (VSVG) strain expressing *F. tularensis* LVS OxyR protein fused with the C-terminal VSVG tag was constructed as described earlier (16). The tagging integration vector, p*KL02*, encoding a *rpoC*-VSVG tag protein, kindly provided by Dr. Simon Dove (Boston Children’s Hospital Division of Infectious Diseases, Boston, MA), was used as a positive control. ChIP assay was performed using three biological replicates as previously described (16). Briefly, *F. tularensis* LVS-*oxyR*-VSVG, and *rpoC*-VSVG tag strains were cultured in 50 mL MH-broth at 37°C with constant shaking. At an OD_600_ of approximately 0.4, rifampicin (50 µg/mL; Sigma) was added to the *rpoC*-VSVG cultures for 30 minutes before crosslinking. All cultures were then crosslinked with 1% formaldehyde for 30 minutes, followed by quenching with 250 mM glycine for 5 minutes. Cells were washed thrice with 1×PBS, resuspended in lysis buffer containing protease inhibitors (Sigma), and sonicated to lyse cells and shear chromosomal DNA to fragments of ∼500 base pairs. Cell debris was removed by centrifugation, and the supernatants were adjusted for salt concentration. Immunoprecipitation was performed overnight at 4°C using anti-VSVG agarose beads (Sigma). A 50 µL aliquot of the supernatant was diluted in 200 µL TE + 1% SDS to be used as an input control. Immunoprecipitates were washed five times with IPP150 buffer, twice with TE buffer, and eluted using 150 µL elution buffer and 100 µL TE + 1% SDS buffer, respectively (16). Elutes and input samples were incubated overnight at 65°C to reverse crosslinking, and the DNA was purified using a PCR purification kit (Qiagen). ChIP and the input DNA samples were analyzed by qPCR to determine the proportion of specific DNA fragments. qPCR values were normalized to inputs using an internal *fopA* coding region control. ChIP assays using wild-type *F. tularensis* LVS (mock) and rpoC-VSVG (positive control) strains were included for comparison. Results were expressed as relative enrichments of the detected fragments. The primer sequences used for qPCR are detailed in **Table 2**.

### Electrophoretic mobility shift assay (EMSA)

The EMSA was performed using bacterial lysates and the LightShift Chemiluminescent EMSA kit (Thermo Scientific). Briefly, the bacterial lysates were prepared from bacterial cultures grown similar to those described above for the gene transcriptional analysis. Promoter DNA probes were generated from *F. tularensis* LVS genomic DNA by PCR using a biotin-labeled 5′ forward primer (Integrated DNA Technologies) and an unlabeled reverse primer. Competitor DNA was amplified using the same unlabeled primer pair, and both probes and competitors were purified with a PCR purification kit (Invitrogen). Primer sequences MP386/MP387 (**Table 2**) for promoter regions of the *trxB* gene were designed to amplify an 182 bp fragment from −138 to +43 relative to the *trxB* open reading frame. Primer sequences MP402/MP403 (**Table 2**) used for the *pmrA* promoter region were designed to amplify a 505 bp upstream intergenic region. Biotin-labeled primers MP386b and MP402b (**Table 2**) were used for probe generation. The binding of the transcriptional regulator PmrA (Sammons-Jackson et al., 2008) to its promoter served as a positive control to validate protein activity in the Δ*oxyR* mutant lysates. Protein extracts were prepared from *F. tularensis* LVS, the Δ*oxyR* mutant, and transcomplemented strains grown to OD_600_ of 0.5 in MH-broth at 37°C. After harvesting and washing, cells were resuspended in TE buffer (10 mM Tris, pH 7.4; 1 mM EDTA, pH 8.0) with protease inhibitors (Sigma). The cells were lysed by sonication, and the soluble protein fraction was separated by centrifugation for 15 minutes (4000×g) at 4°C. The protein concentrations were determined using BioRad reagents. For binding reactions, 1 ng of DNA probe was incubated with 5 µg of total protein in 20 µL reaction buffer following EMSA kit instructions. Competitor DNA (30 ng) was added as indicated. Reactions were loaded onto a 5% TBE non-denaturing gel (BioRad), electrophoresed in 0.5× TBE buffer, and transferred to Hybond-N+ nylon membranes (Amersham). Membranes were crosslinked and probed with streptavidin-HRP conjugates to detect biotin-labeled DNA. A chemiluminescent substrate was used for visualization, and DNA bands were imaged using a Chemidoc XRS system (BioRad).

### Macrophage invasion assay

To evaluate the intracellular survival of wild-type *F. tularensis* LVS, the *ΔtrxB* mutant, and the transcomplemented strain, a gentamicin protection assay was conducted as described earlier (53). Briefly, the murine macrophage cell line RAW264.7 or bone marrow-derived macrophages (BMDMs) isolated from C57BL/6 mice were infected with *F. tularensis* LVS, the *ΔtrxB* mutant, and the transcomplemented strains at a multiplicity of infection (MOI) of 100. Two hours post-infection, macrophages were treated with gentamicin (100 µg/mL) for 2 hours to eliminate the extracellular bacteria. The gentamicin-containing medium was then replaced with an antibiotic-free medium, followed by incubation at 37°C in 5% CO_2_. Macrophages were harvested at 4- and 24-hours post-infection and lysed using 0.1% sodium deoxycholate. The lysates were serially diluted in sterile PBS and plated on MH-chocolate agar plates for bacterial enumeration. Results were expressed as Log_10_ CFU/mL.

### Statistical analysis

All data were statistically analyzed using One-way ANOVA followed by Tukey-Kramer Multiple Comparison tests or the student’s t-test and were expressed as means ± SEM or SD. The value of *P*<0.05 was considered statistically significant.

## ACKNOWLEDGEMENT

This work was supported by National Institutes of Health Grants R21AI51277 (CSB) and R15AI107698 (MM). The funders had no role in study design, data collection and analysis, decision to publish, or manuscript preparation. No financial conflicts of interest exist regarding the contents of the manuscript and its authors.

## Notes

### Competing Interest Statement

The authors have declared no competing interest.

