## Supplemental Figure S1 for "Role of Thioredoxin Reductase (TrxB) in Oxidative Stress Response of *Francisella tularensis* Live Vaccine Strain"

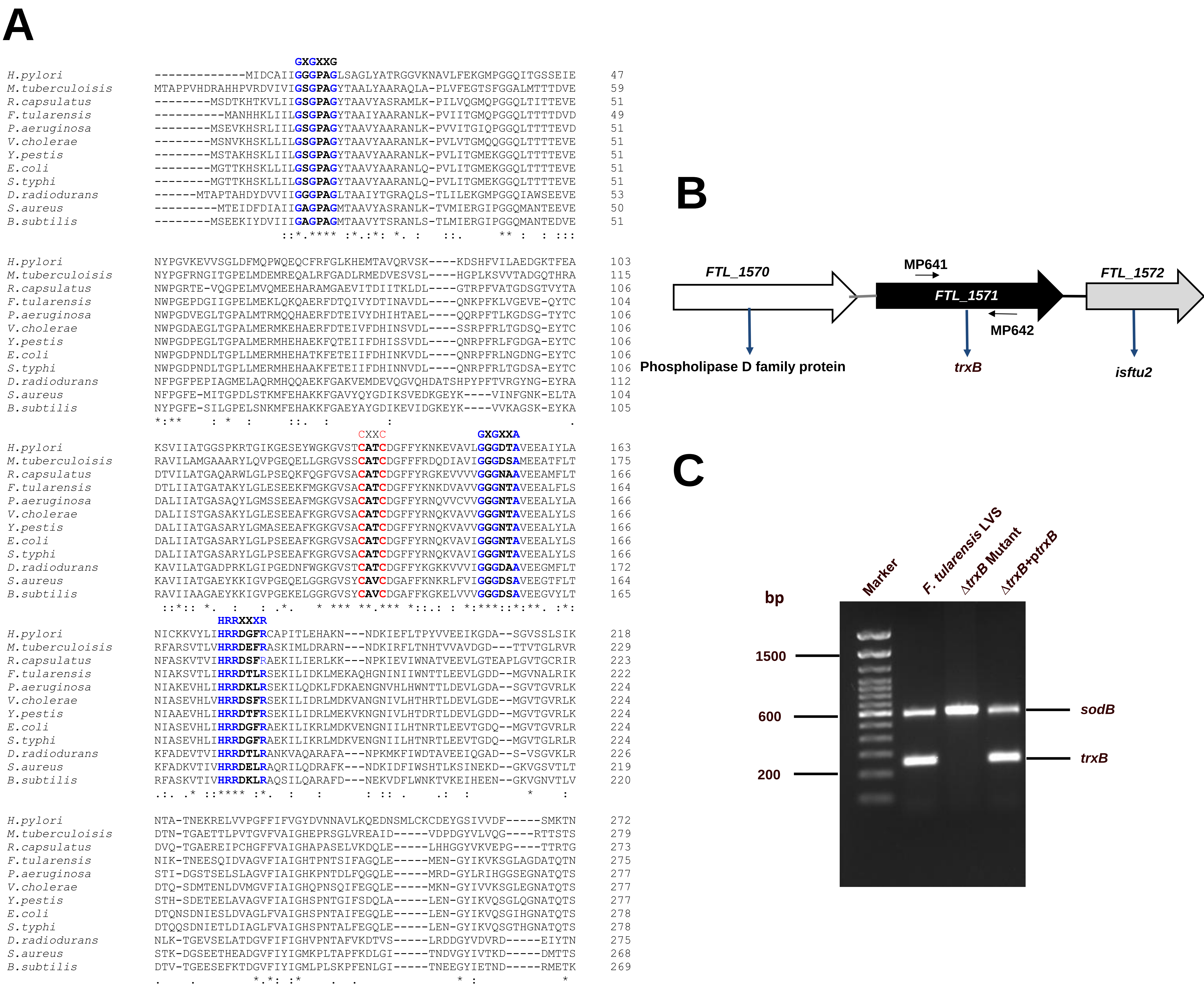


**FIGURE S1: Bioinformatic analysis, Confirmation of *trxB* gene deletion, and transcomplementation. (A)** Multiple amino acid sequence alignments of thioredoxin reductases from selected bacteria were performed using Clustal Omega. Conserved motifs are highlighted in blue or red fonts. The sequences were obtained from the National Center for Biotechnology Information (NCBI) resources. **(B)** Genomic organization of the *trxB* (*FTL_1571*) locus in *F. tularensis* LVS. Arrows indicate the locations of primers used for screening the Δ*trxB* mutant. **(C)** Confirmation of the Δ*trxB* mutant by duplex PCR using *trxB*-specific primers (shown in B) and *sodB* primers as internal controls.
